# Multiple Unfolded Protein Response pathways cooperate to link cytosolic dsDNA release to Stimulator of Interferon Gene (STING) activation

**DOI:** 10.1101/2024.05.10.593557

**Authors:** Tiancheng Hu, Yiping Liu, Jeremy Fleck, Cason King, Elaine Schalk, Zhenyu Zhang, Andrew Mehle, Judith A Smith

## Abstract

The double-stranded DNA (dsDNA) sensor STING has been increasingly implicated in responses to “sterile” endogenous threats and pathogens without nominal DNA or cyclic di-nucleotide stimuli. Previous work showed an endoplasmic reticulum (ER) stress response, known as the unfolded protein response (UPR), activates STING. Herein, we sought to determine if ER stress generated a STING ligand, and to identify the UPR pathways involved. Induction of IFN-β expression following stimulation with the UPR inducer thapsigargin (TPG) or oxygen glucose deprivation required both STING and the dsDNA-sensing cyclic GMP-AMP synthase (cGAS). Furthermore, TPG increased cytosolic mitochondrial DNA, and immunofluorescence visualized dsDNA punctae in murine and human cells, providing a cGAS stimulus. N-acetylcysteine decreased IFN-β induction by TPG, implicating reactive oxygen species (ROS). However, mitoTEMPO, a mitochondrial oxidative stress inhibitor did not impact TPG-induced IFN. On the other hand, inhibiting the inositol requiring enzyme 1 (IRE1) ER stress sensor and its target transcription factor XBP1 decreased the generation of cytosolic dsDNA. iNOS upregulation was XBP1-dependent, and an iNOS inhibitor decreased cytosolic dsDNA and IFN-β, implicating ROS downstream of the IRE1-XBP1 pathway. Inhibition of the PKR-like ER kinase (PERK) pathway also attenuated cytoplasmic dsDNA release. The PERK-regulated apoptotic factor Bim was required for both dsDNA release and IFN-β mRNA induction. Finally, XBP1 and PERK pathways contributed to cytosolic dsDNA release and IFN-induction by the RNA virus, Vesicular Stomatitis Virus (VSV). Together, our findings suggest that ER stressors, including viral pathogens without nominal STING or cGAS ligands such as RNA viruses, trigger multiple canonical UPR pathways that cooperate to activate STING and downstream IFN-β via mitochondrial dsDNA release.

## Introduction

During pathogen invasion, the endoplasmic reticulum (ER)-resident pattern recognition receptor (PRR) Stimulator of Interferon Gene (STING), induces type I interferon (IFN) expression upon binding to cytosolic cyclic di-nucleotides (CDN)^1,2^. CDN ligands derive from bacterial secretion of second messengers, as in the case of *Listeria monocytogenes* produced cyclic-di-AMP^3^. STING also responds indirectly to cytoplasmic DNA via cyclic GMP-AMP synthase (cGAS), a PRR for cytosolic linear double stranded DNA (dsDNA)^4,5^. Upon dsDNA binding, cGAS converts GTP and ATP to the second messenger cyclic GMP-AMP (cGAMP) that potently activates STING^6^. Cytosolic DNA detection is critical in the recognition of viral pathogens, as viruses inject their own DNA into host cells to replicate and lack other pathogen associated molecular patterns (PAMPs) which can be used to detect bacteria, such as flagellin, peptidoglycan, or lipopolysaccharide^7^. Upon activation, STING binds to Tank Binding Kinase 1 (TBK1) and IκB kinase (IKKε), which phosphorylate transcription factors responsible for initiating the innate immune response, such as interferon regulatory factor 3 (IRF3) and nuclear factor κB (NF-κB), ultimately leading to the transcription of *IFNB1* (IFN-β) and other NF-κB dependent inflammatory cytokines^2,8,9^.

Beyond recognition of bacteria and DNA viruses, STING plays a role in a broader array of pathologic conditions, including “sterile” responses to self-DNA, derived from either the nucleus or mitochondria^10-14^. For example, one study showed STING-deficient mice were more resistant to inflammation driven carcinogenesis when exposed to carcinogenic material that released nuclear DNA into the cell^15^. Aberrant mitosis due to DNA damage can drive genomic DNA into the cytosol in “micronuclei”, stimulating cGAS and STING^16,17^. In lupus, oxidized mitochondrial DNA is released into the extracellular milieu, and taken up by bystander cells, leading to STING activation and pathogenic type I IFN production^18,19^. STING also participates in responses to RNA virus infections through unclear mechanisms, though mitochondrial DNA has been implicated in these settings as well^20,21^. Thus, elucidating STING activation is important for understanding inflammation in the broader infectious context and in “sterile” diseases including cancer, cardiovascular disease, kidney diseases, neurodegeneration, aging and autoimmunity^10,13,22-25^.

While previously exploring the relationships between an ER stress response known as the unfolded protein response (UPR), STING, and IFN production, we found that ER stress alone activates innate immune responses as evident by the phosphorylation and nuclear translocation of IRF3, even in the absence of a nominal PRR agonist^26^. Furthermore, the UPR inducer thapsigargin (TPG) induced co-clustering of STING and TBK1 and IRF3 activation was STING-dependent^26^. Interestingly, only some of the methods for inducing ER stress, particularly those that mobilized calcium (ionophore, TPG, and oxygen-glucose deprivation) required STING activation for IRF3 phosphorylation and subsequent IFN-β induction. While ER stress alone induces IFN-β expression at a very low level, ongoing ER stress dramatically synergizes with PRR agonists^26,27^. Since this work, ER stress has been linked to STING activation and inflammation in diverse diseases such as alcoholic liver disease, ischemic brain injury, cardiac hypertrophy, and intracellular bacterial infections^28-31^. Moreover, studies have identified two-way crosstalk between ER stress and STING. In *Listeria* infection, STING was critical for inducing UPR-related “ER-phagy”^32^. During *Brucella* infection, the UPR augments STING induced cytokine production and conversely, STING plays a vital role in enhancing the pathogen induced UPR^33^. Indeed, a “UPR motif” encompassing STING amino acids 322-343 has been identified, which appears critical for STING regulation of calcium homeostasis and associated ER stress^34^. Although these studies have supported the idea of UPR-STING crosstalk, the exact mechanism(s) by which ER stress activates STING remains unclear.

The UPR encompasses three canonical signaling pathways stemming from the activation of ER-stress sensors inositol requiring enzyme 1 (IRE1), activating transcription factor 6 (ATF6) and PKR-like ER kinase (PERK)^35,36^. IRE1 is both a kinase and endonuclease that splices X-box binding protein 1 (XBP1) mRNA, thus generating the active XBP1 transcription factor. PERK activation transiently decreases protein translation. Together these pathways enhance ER function and capacity through new gene transcription and decrease protein load to cope with stress. If stress persists, the UPR initiates apoptosis. Both IRE1 and PERK pathways also regulate re-dox status in the cell through multiple mechanisms^37^. For instance, the inducible nitric oxide synthase (iNOS) gene is an XBP1 transcriptional target. PERK induces oxidating folding chaperones and anti-oxidant responses^38^. Previous work had implicated XBP1 in promoting IFN-β expression^27,26^. Sen et al. identified PERK-dependent pathways regulating type I IFN production during traumatic brain injury^29^. On the other hand, the ATF6 pathway was not involved in TPG-induced IRF3 activation or IFN-β induction^26^. It is not clear how these different UPR pathways impact STING activation.

In this study, we investigated the mechanism of ER stress-dependent STING activation, first determining if ER stress generated nominal cGAS/STING ligands (cytoplasmic dsDNA). We further interrogated the roles of IRE1 and PERK-dependent pathways in regulating cytoplasmic dsDNA release and ER-stress-dependent IFN-β induction. Finally, we determined the effect of IRE1 and PERK inhibition on RNA virus (vesciular stomatitis virus, VSV) induced IFN-β and dsDNA release. Together, our results suggest these two ER stress pathways cooperate to generate cytosolic dsDNA, likely mitochondrial in origin, that triggers cGAS/STING activation.

## Materials and Methods

### Cell culture and treatment

Human HeLa H1 cells (American Type Culture Collection) were maintained in DMEM/high glucose (Mediatech, Manassas, VA, USA) with 10% FBS and antibiotic-antimycotic solution. Murine bone marrow derived macrophages were immortalized with *V-raf/V-myc* as previously described^39^ and maintained in 10% FBS (Hyclone, Logan, Utah USA ) and antibiotic-antimycotic solution (Mediatech) supplemented RPMI 1640 media (Mediatech). Macrophages were isolated from bone marrows of Bim (*Bcl2l11)*^-/-^ mice (gift of Christine Sorenson), STING (*Tmem173*)^-/-^, *Cgas*^-/-^ mice (gifts of Sergio Costa Oliveira), STING mutant Golden Ticket mice (*Tmem173*^gt^, gift from Russel Vance^40^), CHOP (*Ddit3*)^-/-^ (Jackson), or C57BL/6 mice (Jackson). A549 *MAVS*^-/-^ cells^41^ (gift from Craig McCormick) were generated using CRISPR/Cas9 and maintained in DMEM.

To induce ER stress, H1 HeLa cells were treated with 1μM thapsigargin (TPG) or cultured in OGD (oxygen glucose deprivation) conditions: glucose-free DMEM in a hypoxic incubator filled with mixed gas containing 1% O_2_, 5% CO_2_, and 94% N_2_ at 37 °C. For synergy experiments, immortalized macrophages (iMacs) were cultured with 1 μM TPG for one hour and then 10ng/mL lipopolysaccharide (LPS) for a further 3h or poly I:C for another 6h prior to harvest. For MitoTEMPO experiments, iMacs were treated 1h with MitoTEMPO (1, 2 or 5 μM) and then a further 3h with TPG. N-acetylcysteine was added at 0.1 mM for 30 minutes prior to TPG. Thapsigargin (cat #T9033), LPS (cat#L7770), MitoTEMPO (Cat#SML0737), N-Acetyl-L-cysteine (cat# A9165), PERK Inhibitor I, GSK2606414 (cat# 516535), and the IRE1 inhibitor III, 4μ8C (cat#412512) were purchased from Sigma; poly I:C was from Invivogen (31852-29-6); 1400W dihydrochloride (cat # 1415) was purchased from R&D; MitoTracker™ Red CMXRos (cat # M7512) was from Thermofisher.

### Transfection with siRNA

HeLa cells were transfected with 200 pM scrambled control or target siRNA (Thermofisher) using Lipofectamine RNAiMax (Invitrogen) according to the manufacturer’s instructions. Assays were performed at least 24h post-transfection.

### Quantitative PCR

Cells were lysed with TRIzol (Invitrogen) and processed as previously described. Briefly, RNA was extracted with chloroform and precipitated with isopropanol. RNA quality (260/280 ratio) and quantity were assessed by NanoDrop 1000 (Thermo Scientific, Wilmington, DE, USA). Total RNA was treated with DNase I (Invitrogen) prior to reverse transcription using a superscript kit (Invitrogen). Gene expression in cDNA was quantitated by SYBR Green (Invitrogen) fluorescence detected on a StepOne real time PCR system (Thermofisher). Reaction efficiency was assessed using a serially diluted standard curve and product by melting curve. Relative mRNA expression was normalized to the 18S rRNA housekeeping gene using the standard ΔCt/ ΔCt method. Primers were designed using Beacon design software (Premier Biosoft, Palo Alto, CA, USA) or identified from the literature: h*18S* rRNA, F: GGACACGGACAGGATTGACAG3; R: ATCGCTCCACCAACTAAGAACG h *IFNB1*, F: TGGCTAATGT CTATCATCA; R: CTTCAGTTTCGGAGGTAA h *NOS2*, F: CCTGGTACGGGCATTGCTCC, R: GCTCATGCGGCCTCCTTTGA m *18S* F: GGACACGGACAGGATTGACAG; R: ATCGCTCCACCAACTAAGAACG m *Ifnb1* F: ACTAGAGGAAAAGCAAGAGGAAAG; R: CCACCATCCAGGCGTAGC h *XBP1* F: TAGTGTCTAAGGAATGAT; R: CCAGTAATATGTCTCAATA m *Xbp1*(s): F: GAGTCCGCAGCAGGTG; R:GTGTCAGAGTCCTCCATGGGA VSV^42^ P/M intergenic region: 5’-TCCTGCTCGGCCTGAGATAC-3’ VSG M/G intergenic region: 5’-TCCTGGATTCTATCAGCCACTT-3’

### Mitochondrial (mt) DNA detection^43,44^

MtDNA was isolated and quantified using standard methods^43^. Briefly, cell pellets were resuspended in lysis buffer with 150 mM NaCl, 50mM HEPES pH7.4 and 20μg/mL digitonin. After 10 min, samples were spun down in a refrigerated microfuge at 1000g for 10 min. Supernatants were transferred to fresh tubes and centrifuged another 10 min at 17000g. DNA in these supernatants was concentrated and cleaned with a kit (Zymo). Whole cell lysates from matched samples were analyzed in parallel for normalization. MtDNA was isolated using Dneasy Blood and Tissue kits (Qiagen). qPCR was perfomed on these cytosolic supernatants and whole cell extracts using the following mtDNA primers: F: CCTAGGGATAACAG GCAAT; R: 5: TAGAAGAGCGATGGTGAGAG^45,46^. Cytosolic mtDNA was normalized to whole cell extracts, and the value in the vehicle control was set to 1.

### Immunofluorescence studies

Cells were plated on coverslips in 60-mm dishes for 24h prior to treatment. After treatment, cells were washed with PBS and then fixed in 4% paraformaldehyde 30 minutes (min) at room temperature. Cells were then washed with PBS, Tris A buffer (0.1 M [pH 7.6] Tris and 0.1% Triton X-100), and Tris B buffer (0.1 M [pH 7.6] Tris, 0.1% Triton X-100, and 0.2% BSA) 3x 5 min each and incubated with 10% goat serum in Tris B buffer for 1h. Primary antibodies (Ab) were added in Tris B buffer and cells incubated at 4°C overnight. After washing the cells with Tris A 3x 5 min, secondary fluorescence-conjugated Ab was added and samples incubated 1h at room temperature. Cells were washed with PBS 3x 5 min and the coverslips mounted on slides with ProLong Gold antifade reagent with DAPI nuclear stain (Invitrogen). For negative controls, the same concentration of primary mouse IgG (Sigma-Aldrich) or rabbit IgG (Sigma-Aldrich) was added. Images were acquired on a Nikon A1Rs confocal fluorescent microscope (Nikon). Primary antibodies: STING (D2P2F) Rabbit mAb (Cat#13647 Cell Signaling Technology-CST, Danvers, MA, USA), dsDNA mouse mAb (cat# sc-58749 Santa Cruz biotechnology, Santa Cruz, CA USA); mouse anti-RVC VP1 (ref); MitoTracker™ Red CMXRos (cat#M7512, Thermofisher scientific, USA). Secondary antibodies: rabbit anti-mouse IgG Alexa Fluor 488, or rabbit anti-mouse IgG Alexa Fluor 594 were from Thermofisher.

### Quantification of cytosolic dsDNA

Fluorescence microscopy images were analyzed using Image J to remove the nuclear DNA. Original and nucleus-free images were then edited in ImageJ by changing the threshold color setting to B&W and analyzed through the “Analyze Particles” tool to identify spots with a size of 0-100 and a circularity of 0.00-1.00, ensuring measurement of only cytoplasmic dsDNA. Cytosolic dsDNA “density” = proportion of total fluorescence accounted for by speckles ((Total fluorescence – nuclei)/cell number per field).

### Statistics

Multiple sample comparisons were made with ANOVA and two-way comparisons were performed using Student’s T-test. Bars represent mean expression of one representative experiment in triplicate, or means of at least 3 independent experiments, and error bars denote STD/SEM accordingly. In Figures *p<0.05, **p<0.01, ***p<0.005, ****p<0.001.

### Vesicular stomatitis virus (VSV) infection

Vesicular stomatitis Indiana virus carrying GFP was described previously^42^. Virus was diluted in OptiMEM supplemented with 0.2% BSA and 1x Pen/Strep and applied to HeLa or A549 cells at an MOI=1. Following inoculation, cells were grown in DMEM supplemented with 10% FBS for 4h prior to fixation with 4% paraformalin to visualize dsDNA. IFN-β mRNA expression was detected after 6h and protein in viral supernatants at 8h using an R&D ELISA kit (cat#DIFNB0).

## Results

### ER stress induced IFN-β mRNA requires cGAS and STING

Previous work had shown ER stress induces STING clustering and the association of STING with TBK1^26^. However, it was not known whether ER stress activates STING directly, in the absence of self-ligand, or via a cGAS-generated intermediate. Treating immortalized macrophage (iMac) cells with TPG, which inhibits SERCA pump function and depletes ER calcium stores,^47^ significantly induced IFN-β mRNA expression (Figure 1A) and promoted STING clustering (Figure 1B)^8,48^. To determine if STING and cGAS were required for TPG-induced IFN-β (*Ifnb1*) mRNA, we used immortalized macrophages (iMacs) derived from *Tmem173*^-/-^ and *Cgas*^-/-^ mice. IFN-β mRNA induction was completely abrogated in *Cgas*^-/-^ and *Tmem173*^-/-^ macrophages (Figure 1C). These requirements were then tested in another model of ER stress, oxygen glucose deprivation (OGD). In contrast to wild type (WT) macrophages (Figure 1D), *Tmem173*^-/-^ and *Cgas*^-/-^ macrophages were unable to upregulate *Ifnb1* transcripts during OGD. It was possible, given the known effects of STING on the UPR, that the profound inhibition with cGAS or STING deficiency simply reflected an abrogated UPR. Preliminary data (Figure S1) suggests this is not the case. ER stress induces very low levels of IFN, but synergizes dramatically with PRR agonists such as lipopolysaccharide (LPS) to upregulate IFN expression and protein secretion^26,27^. Both cGAS and STING were required for full ER stress-dependent synergism with LPS (Figure 1E), consistent with previously reported STING knockdown data^26^. In contrast, Poly I:C-TPG synergism did not require cGAS and STING (Figure S2), possibly reflecting alternative mechanisms of synergy. The requirement for cGAS in TPG and OGD induced IFN induction supports the idea that ER stress activates STING via a cytosolic dsDNA intermediary, rather than directly.

**Figure 1:**
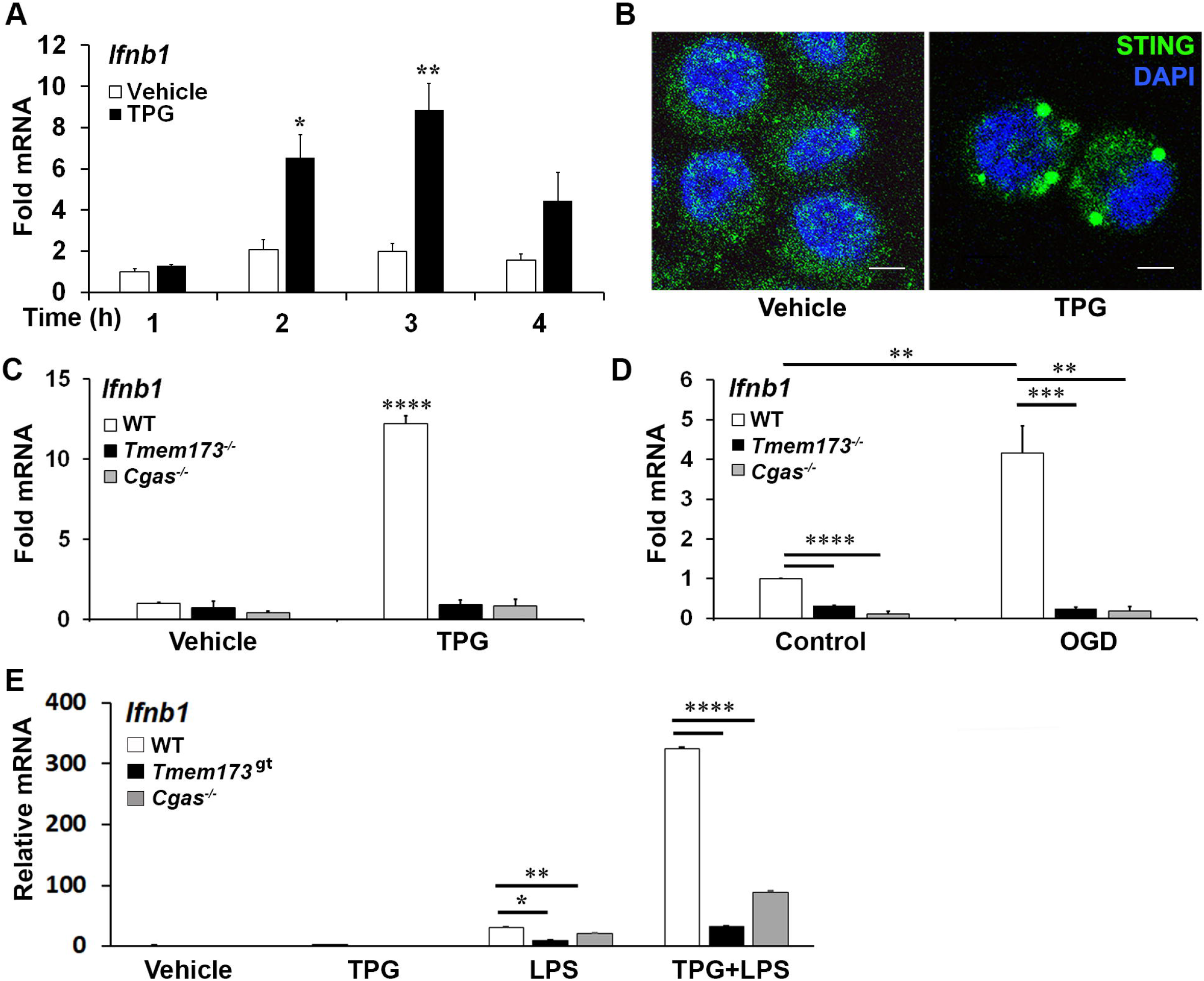
ER stress-dependent *Ifnb1* (IFN-β) mRNA expression during TPG treatment and oxygen glucose deprivation (OGD) requires both STING and cGAS. (A) iMacs were treated with 1μM TPG (black bars) or DMSO vehicle control (open bars) for the specified times. *Ifnb1* mRNA expression was determined by quantitative (q) PCR, normalized to 18S RNA and to vehicle treated control (set=1). *P<0.05 and **p<0.01 comparing TPG and DMSO samples within time points. (B) HeLa cells were treated with 1μM TPG for 3h, fixed, and incubated with anti-STING followed by anti-mouse IgG Alexa Fluor 488 (green), and then stained with DAPI (blue nuclei). Visualization was by immunofluorescence microscopy. Scale bars are 5 μm(C) Wild type (WT, open bars) STING (*Tmem173*)^-/-^ (black bars) and *Cgas*^-/-^ (gray bars) iMacs were treated with 1μM TPG for 3h and *Ifnb1* mRNA quantified as above. ****P<0.001 in treated WT vs. all knockouts and vehicle control. RNA levels were normalized to vehicle treated WT control (set=1) (D) iMacs were subject to oxygen glucose deprivation (OGD) for 2h and IFN-β expression was quantitated with qPCR as above, with fold RNA vs WT control. ****p<0.001, ***p<0.005 and **p<0.01. (E) WT, *Cgas*^-/-^ or STING null mutant (Golden Ticket, *Tmem173*^*gt*^) macrophages were stimulated with 1μM TPG for 1h and then 10ng/mL LPS for 3h prior to harvest for RNA. RNA levels were normalized to 18S RNA. Bars represent means of 2-3 (E), 3 (A, D) or 5 (C) independent experiments and errors are SEM.

### ER stress results in the release of cytosolic dsDNA

The TPG model of ER stress is “sterile”, performed in the absence of nominal pathogens or PAMPs, suggesting that the DNA ligand activating cGAS and STING is endogenous. Whereas dsDNA appeared primarily restricted to the nucleus using immunofluorescence microscopy in control settings, following TPG stimulation dsDNA was evident in small clusters (green speckles) throughout the cytoplasm, outside of the nucleus in HeLa and iMac cells (Figure 2A). The close spatial and functional relationships between the ER and mitochondria suggested mitochondria might be sensitive to ER stress and thus a source of the DNA speckles^22,25^. To determine if the extra-nuclear puncta of dsDNA co-localized with mitochondria, we used MitoTracker dye. In TPG, but not vehicle treated cells, mitochondrial staining overlapped with the small dsDNA speckles (yellow, Figure 2B). Also, the contours of the nucleus still appeared intact in all experiments, without evident micronuclei. Furthermore, quantitative PCR of cytosolic extracts confirmed an increase in mitochondrial DNA following TPG (Figure 2C), which correlated well with the dsDNA cytoplasmic immunofluorescence signals. Previous work using this dsDNA antibody also showed a correlation of the fluorescent punctae with increased cytosolic mitochondrial DNA and DNAse sensitive ISRE-reporter stimulating cytoplasmic nucleic acid^44^. Together, these results suggest the endogenous dsDNA release stimulated by ER stress is mitochondrial in origin, rather than nuclear.

**Figure 2.**
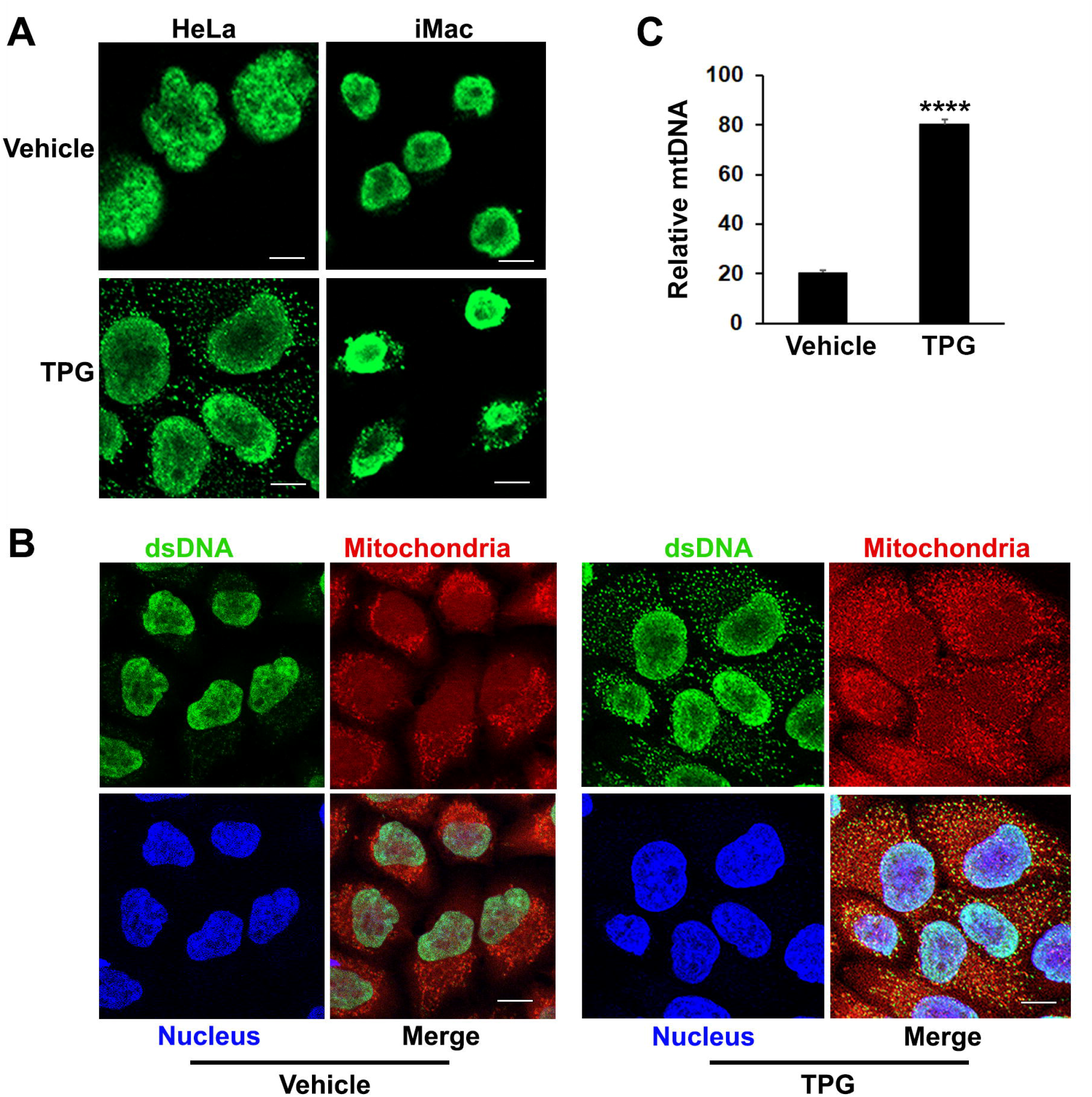
TPG treatment results in the release of cytoplasmic dsDNA. (A) HeLa cells (left) or iMacs (right) were treated with 1μM TPG or DMSO vehicle control for 1h. Cells were then fixed and incubated with anti-dsDNA followed by anti-mouse Alexa Fluor 488 antibodies. (B) HeLa cells were stained with MitoTracker for 30 min and treated with DMSO vehicle control (left panels) or 1μM TPG (right panels) for 1h. Fixed cells were stained with DAPI (nuclei) and anti-dsDNA as above. Visualization was by immunofluorescence microscopy. Scale bars are 10 μm (C) HeLa cells were treated with DMSO (vehicle) or TPG for 1h as above. Mitochondrial DNA (mtDNA) was quantitated by qPCR with normalization of the cytosolic fraction to whole cell extract. N=2 and p<0.001.

### The role of the IRE1 axis of the UPR in cytosolic dsDNA release

In addition to the canonical UPR signaling cascades, ER stress causes derangements in intracellular calcium and generates oxidative stress, which may impact mitochondrial stability. To determine if oxidative stress contributes to ER-stress induced IFN-β, we pre-treated cells with the antioxidant N-acetylcysteine (NAC). Indeed, NAC significantly reduced induction of IFN-β mRNA by TPG (Figure 3A).

**Figure 3.**
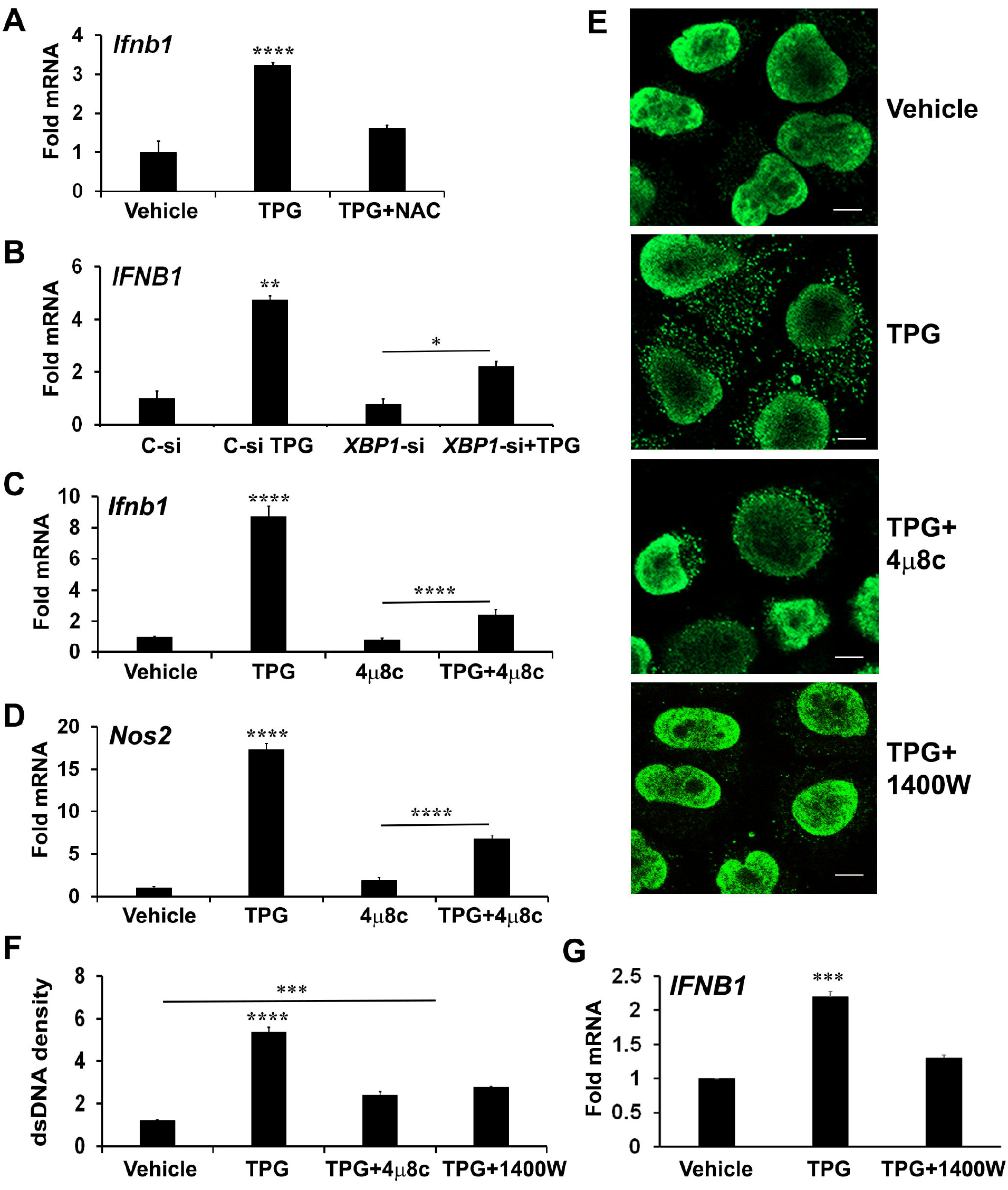
XBP1 and iNOS contribute to UPR induced cytosolic dsDNA release. (A) iMacs were treated with DMSO vehicle control only or pre-treated with 0.1 mM N-acetylcysteine (NAC) for 30 min and then 1μM of TPG for 3h. IFN-β expression was quantitated using qPCR (Fold RNA) with vehicle set=1. Results are from N=3 experiments. **** P-values are in comparison to vehicle or TPG+NAC treatments. (B) HeLa cells were transfected with siRNA targeting *XBP1* (*XBP1*-si) or scrambled control (C-si). 24h later cells were stimulated with TPG for 3h. qPCR determined expression levels of *IFNB1* mRNA. N=2 experiments. **p<0.01 vs. all other conditions, *p<0.05. (C, D) iMacs were treated with pre-treated with 10 μM 4μ8C 30 min then 1μM TPG for 3h as indicated and IFNβ (*Ifnb1*) mRNA (C) or iNOS (*Nos2*) mRNA (D) quantitated as above. N=4 experiments. ****p<0.001 comparing TPG with all other conditions and TPG+4μ8c with 4μ8c. (E and F) HeLa cells were pre-treated with 4μ8C or 1μM 1400W followed by 1h TPG as indicated, then fixed and incubated with anti-dsDNA and anti-mouse IgG Alexa Fluor 488 antibodies. dsDNA particles were imaged using confocal microscopy (E) and cytosolic fluorescence quantitated using image J. Scale bars are 10 μm (F). ****p<0.001 comparing TPG alone with other conditions, ***p<0.005 vs DMSO vehicle control. Results are from 3 fields in one experiment and representative of N=2. G) HeLa cells were pre-treated with 1400W for 1h followed by TPG for 3h. RNA was processed for qPCR and analyzed for *IFNB1* expression. N=3. ***p<0.005 vs vehicle only or 1400W+TPG treatment.

Reactive oxygen and nitrogen species derive from multiple cellular locations including the ER, cytosol, and mitochondria. To determine if mitochondrial ROS contributed to TPG-induced IFN, we pre-treated cells with MitoTEMPO, which suppresses mitochondrial ROS^19^. However, MitoTEMPO (1-5 μM) had no effect on TPG-induced IFN-β mRNA induction (Figure S2), suggesting mitochondrial ROS are not a major source of STING activation in our experimental system.

Therefore, we further investigated a canonical UPR pathway (the IRE1-XBP1 pathway) previously implicated in IFN induction and re-dox regulation^27,49^. We confirmed that inhibiting the XBP1 transcription factor with either small interfering RNA (siRNA) or the IRE1 endonuclease inhibitor 4μ8c (Figure S3) blocked ER stress-induced *IFNB1* expression (Figure 3B, 3C)^27,50^. The addition of 4μ8C also reduced the number of TPG-induced dsDNA cytoplasmic clusters (Figure 3E). These results suggested XBP1 contributed to ER stress-induced dsDNA release. 4μ8c significantly attenuated iNOS induction, as expected, supporting the requirement for XBP1 upstream of iNOS^38^. (Figure 3D). To determine if XBP1-dependent iNOS was required for dsDNA release, we used 1400W, an iNOS inhibitor^51^. A significant decrease in both IFN-β expression and cytosolic dsDNA was observed with 1400W pre-treatment as compared to TPG treatment alone (Figure 3E-G).

### The PERK pathway and downstream Bim contribute to ER stress-mediated dsDNA release

Treatment with 4μ8C or NAC only partially inhibited dsDNA release, prompting further investigation into the role of another UPR branch, the PERK pathway. During chronic ER stress, PERK leads to apoptosis in part through the induction of C/EBP homologous protein (CHOP) and CHOP-dependent Bim expression. Bim, in turn, regulates Bax-Bak dependent changes in mitochondrial permeability^52-54^. Bax-Bak mitochondrial pores may allow for mitochondrial DNA release into the cytoplasm^53,55^. Treating HeLa cells with the PERK inhibitor I (GSK2606414, Figure S4) significantly reduced TPG-induced IFN-β mRNA expression (Figure 4A) and decreased cytosolic mtDNA (Figure 4B). Whereas both 4μ8C and PERK I attenuated dsDNA release to a similar extent, the combination decreased the cytosolic dsDNA back down to vehicle control levels (Figure 4C and 4D). The PERK I inhibitor also decreased TPG-induced Bim (*BCL2L11*) mRNA (Figure 4E). TPG treatment of Bim (*Bcl2l11)*^-/-^ murine macrophages elicited less cytosolic dsDNA release (Figure 4F) and abrogated *Ifnb1* up regulation (Figure 4G), suggesting a critical role for Bim in ER stress-dependent IFN-β induction.

**Figure 4.**
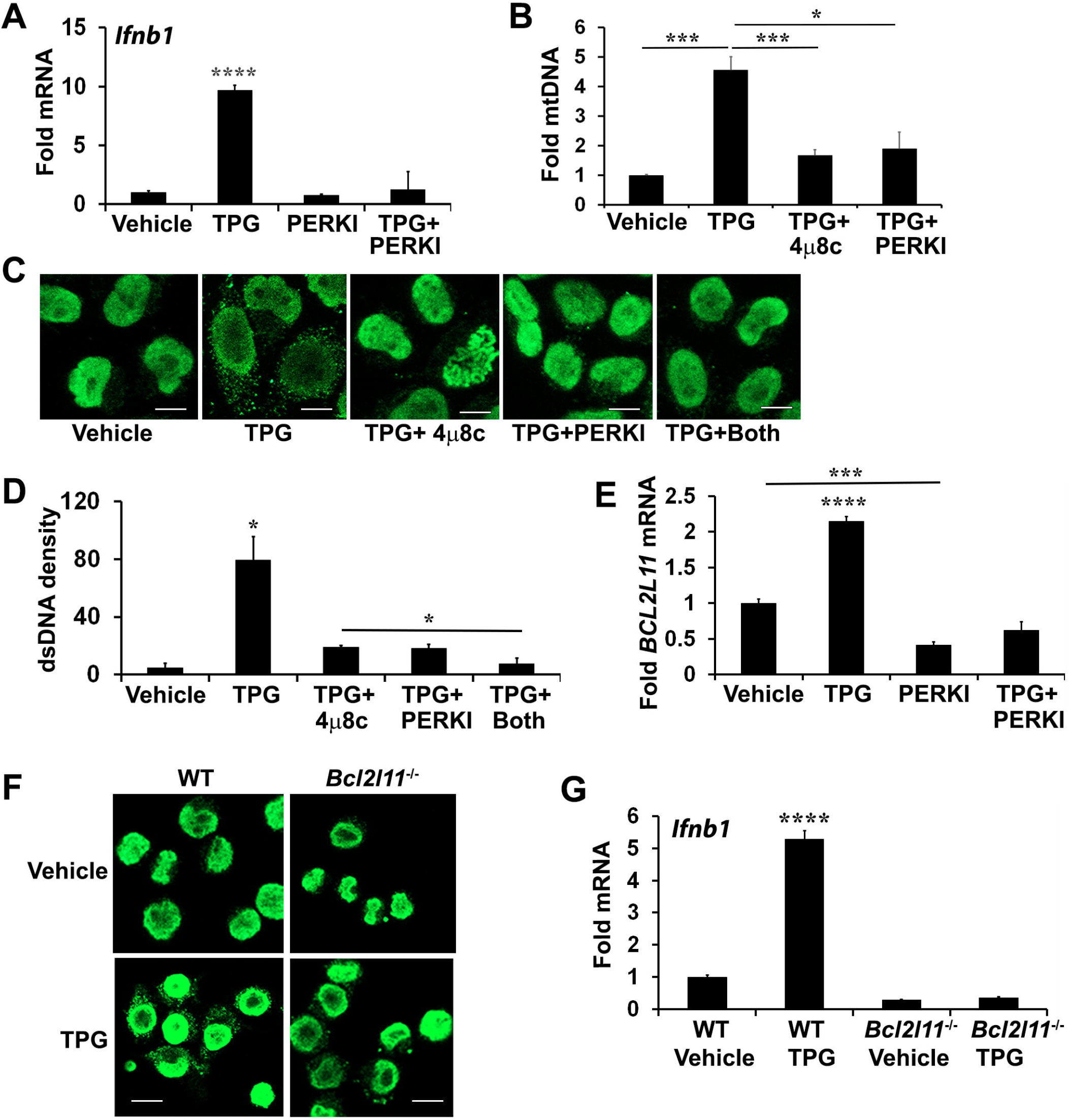
PERK and Bim regulate TPG-dependent *IFNB1* induction and cytoplasmic dsDNA release. (A) iMac cells were pre-treated with 10μM of PERK I (GSK2606414) for 30 min before stimulation with DMSO vehicle or 1μM of TPG for 3h. *Ifnb1* mRNA was quantitated using qPCR. N=3 exp and ****p<0.001 comparing TPG with other conditions. Fold RNA is vs. vehicle control. (B) HeLa cells were pre-treated with 4μ8c or PERKI prior to stimulation with TPG for 1h. mtDNA in cytosolic fractions were quantitated by qPCR and normalized to mtDNA in whole cell extracts. N=3. *p<0.05 and ***p<0.005 in pairwise comparisons. (C) HeLa cells were treated as in (B) and dsDNA was visualized using immunofluorescence. (D) The dsDNA fluorescence densities were quantitated with Image J. Results are averages from 3 fields and are representative of 3 independent experiments. *p<0.05 comparing TPG with all other conditions and TPG+4μ8c with TPG+Both (E) HeLa cells were treated as in (A) and Bim (*BCL2L11*) mRNA quantitated by qPCR as above. ****p<0.001 for TPG vs. all other conditions and ***p<0.005 comparing vehicle only vs PERK I treated cells. (F) WT or Bim (*Bcl2l11*)^-/-^ iMacs were stimulated with 1 μM TPG for 1h prior to fixation and dsDNA staining for immunoflourescence. Scale bars are 10 μm. (G) or 3h prior to harvesting for RNA extraction and qPCR. ****P-value compares TPG treated WT iMacs and either untreated cells or TPG-stimulated *Bcl2l11*^-/-^ cells. N=3.

### ER stress pathways regulate cytosolic dsDNA release and IFN-β production during vesicular stomatitus virus (VSV) infection

Several studies have shown that RNA viruses, which lack nominal STING/cGAS ligands, cause mitochondrial damage and release of mtDNA^56,57^. In this regard, we recently showed influenza A virus triggers increases in cytosolic mtDNA^44^. Moreover, some RNA viruses such as VSV, Dengue virus and West Nile virus are restricted by STING expression and many have developed mechanisms to counteract STING^21,58,59^. Evidence also supports STING-RNA sensing PRR crosstalk^60^. Although viral mediators have been identified that threaten mitochondrial integrity and trigger cytosolic DNA release, the role of UPR pathways in virus-dependent cytosolic dsDNA release are unknown^43,61,62^.

We investigated UPR-STING pathways in the context of RNA viral infection using an *in vitro* VSV infection model. STING has been reported to restrict VSV replication in murine embryonic fibroblasts (MEFs) and in mice^8,58^. We confirmed that in STING mutant macrophages, VSV replicates to a greater extent (Figure 5A). Reciprocally, VSV induced IFN-β expression was decreased in *Tmem173*^-/-^ macrophages (Figure 5B). To test the hypothesis that VSV elicits a STING-stimulating agonist beyond RNA, we used MAVS^-/-^ bronchial epithelial cells (Figure 5C), which revealed residual *IFNB1* mRNA induction of at least one log. As previously seen with influenza^13^, VSV also triggered an increase in anti-dsDNA antibody detected cytosolic punctae in infected cultures (Figure 5D, top 2 rows). Interestingly, dsDNA speckles were evident in some cells where VSV-GFP was not detected.

**Figure 5.**
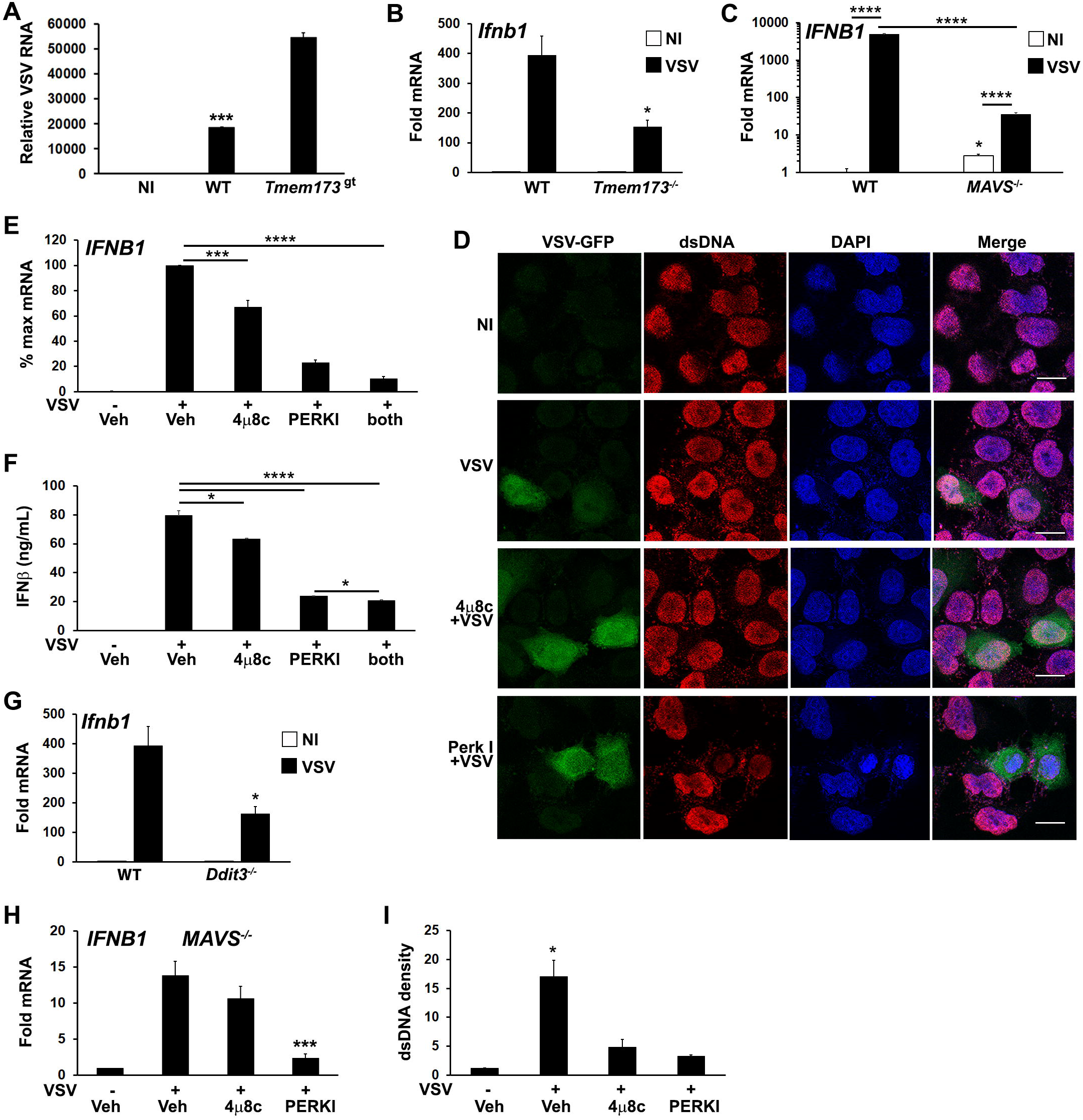
Both PERK and IRE1 UPR pathways regulate VSV-induced dsDNA release and *IFNB1* induction. A) Wild type (WT) or STING mutant (*Tmem173*^gt^) iMacs were infected with VSV (MOI=1) for 24h, harvested for RNA and VSV detected by qPCR^42^ with normalization to 18S rRNA. NI=non-infected WT iMacs. *** p<0.005 for WT vs NI and infected STING mutant cells. N=2 independent experiments. B) WT or *Tmem173*^-/-^ iMacs were uninfected or infected with VSV for 6h prior to harvest for mRNA and *Ifnb1* expression quantifed and normalized as above. * P<0.05 comparing WT and *Tmem173*^-/-^ infected macrophages. N=3. C) *MAVS*^-/-^ A549 cells were uninfected (NI, open bars) or infected (black bars) with VSV for 6h prior to harvest for mRNA, and *IFNB1* expression quantified as above. N=4, with fold change vs uninfected WT set=1. ****p<0.001, *p<0.05 vs WT NI. N=3. D) HeLa cells were pretreated with vehicle or UPR inhibitors and then infected with VSV-GFP for 4h. Cells were fixed and examined for VSV and dsDNA using immunofluorescence microscopy. A rabbit anti mouse Alexa-fluor 594 secondary antibody was used to visualize dsDNA. Results are representative of 3 independent experiments. Scale bars are 10 μm E) A549 cells were pre-treated with vehicle (Veh), the IRE1 inhibitor 4μ8c, or the PERK inhibitor (PERKI) then infected with VSV for 6h prior to harvest for RNA. *IFNB1* expression was quantified by qPCR with normalization to 18S rRNA and VSV infected vehicle treated control (set=100% for maximum mRNA). N=7. ***p<0.005 for VSV infected DMSO vs 4μ8c and ****p<0.001 for all other pairwise comparisons. F) A549 cells were treated and infected as in (E) for 8h, and then IFNβ in supernatants quantified using ELISA. *p<0.05, ****p<0.001. N=3. G) WT or CHOP (*Ddit3*) iMacs were infected with VSV for 6h prior to harvest for mRNA and *Ifnb1* expression quantified by qPCR as above. *p<0.05 vs WT. N=3. H) *MAVS*^-/-^ A549 bronchial cells were pretreated with DMSO vehicle control (Veh), 4μ8c or PERKI for 30 minutes, then infected with VSV for 6 hours. *IFNB1* mRNA was quantitated by qPCR with normalization to 18S rRNA and non-infected vehicle treated control (set=1). Results are from N=3 and ***p<0.005 vs. VSV infected vehicle control. I) The cytoplasmic dsDNA fluorescence densities from (D) were quantitated with Image J. Results are averages from 3 fields and are representative of 3 independent experiments. *p<0.05 comparing vehicle pre-treated VSV with UPR inhibitors and with uninfected cells.

To further examine the roles of IRE1 and PERK pathways in IFN induction and cytsolic dsDNA release, A549 bronchial cells were pre-treated with the 4μ8c and PERK inhibitors prior to VSV infection. Treatment with these inhibitors reduced *IFNB1* expression in A549 and HeLa cells following infection (Figures 5E, S5), with the PERK inhibitor exhibiting a greater effect. *IFNB1* mRNA expression correlated with secreted IFN-β protein (Figure 5F). Consistent with these results, VSV induction of *Ifnb1* mRNA was also decreased in CHOP (*Ddit3)*^-/-^ cells (Figure 5G). *MAVS*-/- cells also displayed decreased IFN expression in the presence of UPR inhibitors, supporting an RNA sensing-independent effect of UPR pathway inhibition (Figure 5H). Treatment with 4μ8c and PERK I also decreased dsDNA release from VSV treated cells, (Figure 5D and 5I). The inhibition of cytosolic dsDNA release by UPR inhibitors was consistent with an upstream role generating STING ligands during VSV infection. PERK and IRE1 pathways have pleiotropic effects on viral replication beyond IFN expression, and different viruses both induce and inhibit various UPR signaling pathways to enhance their success^63-67^. In the case of VSV, inhibition of IRE1 produced a net neutral effect and PERK inhibition was deleterious to viral replication (Figure S6).

## Discussion

STING has been increasingly recognized to play a role in non-infectious pathologies such as cancer, autoimmunity and ischemia as well as in infections lacking a nominal STING agonist such as RNA virus infections^22^. However, the mechanisms by which ER stress activates STING in the absence of nominal ligands have been unclear. In this study we have shown specific ER stressors induce IFN-β mRNA in both a STING and cGAS-dependent manner. Furthermore, stimulation of cells with ER stress-inducing pharmacologic agents, OGD, or viral infection resulted in the appearance of abundant dsDNA particles in the cytoplasm, which could serve as agonists to stimulate cGAS, and thus STING. These particles were clearly evident during both pharmacologic UPR and viral infection, despite cytoplasmic DNases, suggesting a robustly induced process. Independent canonical UPR pathways stemming from activation of IRE1 endonuclease/XBP1 and PERK both contributed to dsDNA release (diagram, Figure 6).

**Figure 6.**
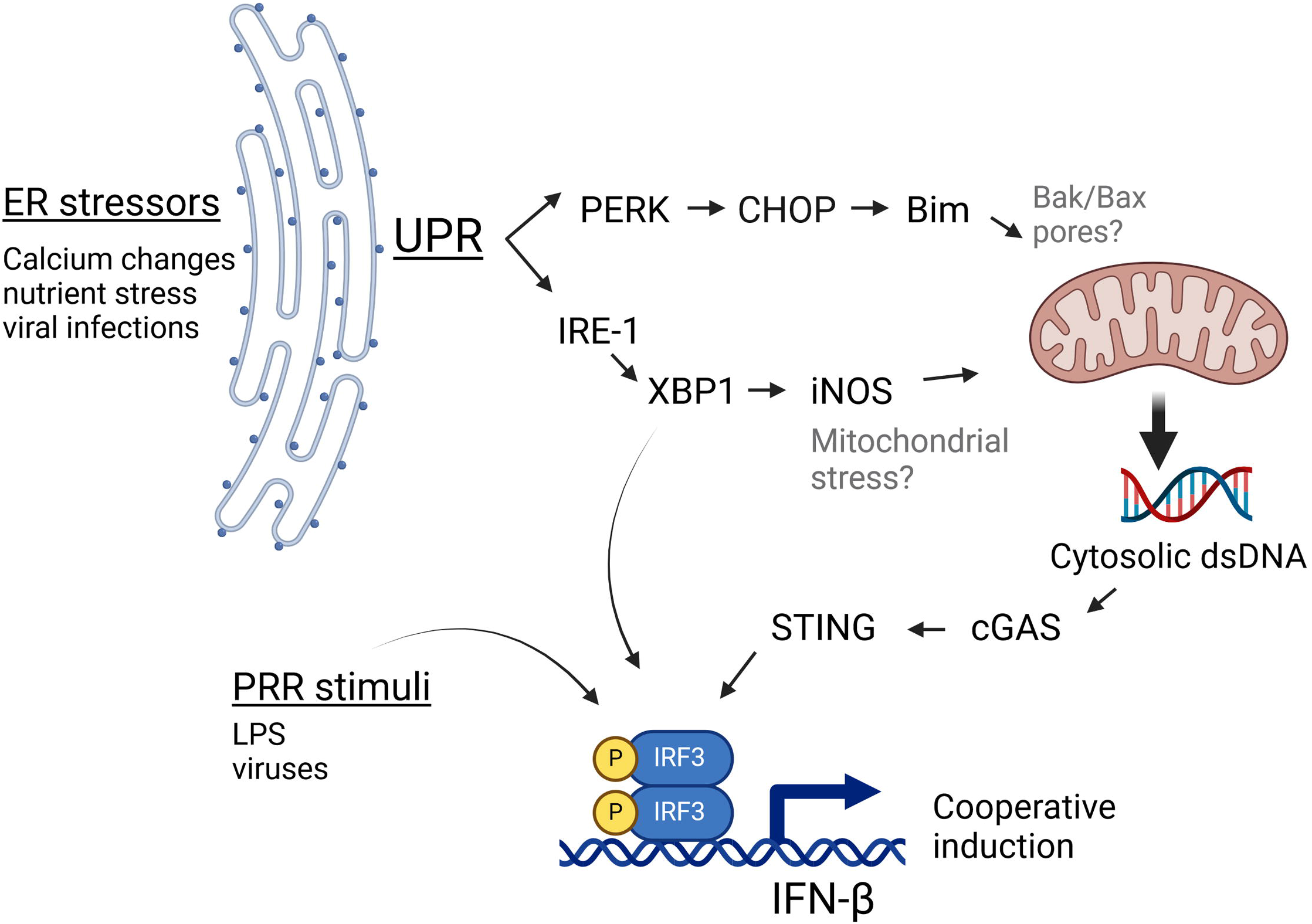
IRE1 and PERK-dependent pathways cooperate to generate cGAS/STING activating dsDNA release. During ER stress, the UPR activates PERK and IRE1 pathways. PERK induces the expression of CHOP and Bim, which influence mitochondrial integrity. IRE1 endonuclease activation results in the translation of the full length XBP1 transcription factor and thus induction of iNOS. Together, these 2 pathways both contribute to the ER stress-induced increase in cytosolic dsDNA, which may serve as a nominal stimulus for cGAS/STING dependent IFN-β induction. Bak/Bax placement within this schematic (in gray) derive from the literature^52^, whereas black font molecules were addressed in this study. This Figure was made using BioRender.

Downstream of XBP1, one pathway involves iNOS induction, though the XBP1-iNOS contribution to dsDNA release and IFN induction appears partial. PERK and Bim also regulated cytoplasmic dsDNA release. Inhibiting both XBP1 and PERK pathways together was additive in suppressing dsDNA release. However, the 2 pathways may also work together. For instance, one might envision that the XBP1-iNOS pathway generates mitochondrial stress, but Bim induction is required for increased dsDNA release from the stressed mitochondria. In this scenario, interfering with either XBP1 or PERK signaling diminishes the dsDNA release. Together the data from these studies suggests that during non-infectious pathologies or infections associated with ER stress, canonical UPR pathways cooperate to stimulate STING-dependent immune responses.

Endogenous DNA from the nuclei or mitochondria can serve as cGAS stimuli that generate cGAMP for STING activation. Nuclear DNA seems to be important in the setting of cancer, where DNA damage causes aberrant mitoses and generation of cytosolic micronuclei^16,17^. In our studies, the nuclei appeared smooth and intact, without obvious micronuclei. Moreover, the rapid time frame of IFN-β mRNA induction (within a few hours) argues against a cell division requirement. Here we showed the ER stressors TPG and OGD, as well as VSV infection, generated numerous cytoplasmic dsDNA speckles. Both the co-localization of cytoplasmic DNA with mitochondria and increase in cytoplasmic mtDNA also suggested these DNA speckles are mitochondrial in origin. It is not clear why the mtDNA was not readily visible prior to stimulation; the antigen recognized by our dsDNA antibody may require dissociation from genome packaging proteins present in intact mitochondria such as mitochondrial transcription factor A (TFAM)^18,68,69^. Another possibility is that cytosolic release results in clumping or clustering of the tiny mitochondrial genomic material.

The partial effect of XBP1 inhibition prompted examination of the other UPR arms. In other studies, the PERK pathway plays both positive and negative roles in regulating type I IFN, depending upon context^29,70-73^. As an example, during infection, PERK may antagonize IFN sensing by supporting IFNAR1 degradation^74^. In our in vitro models, PERK-dependent pathways supported cytosolic dsDNA release and IFN induction. However, the net effect of PERK inhibition in *suppressing* VSV replication suggests multiple roles for the PERK pathway during infection. Previous work had implicated PERK in ER stress-regulated IFN-β in dendritic cells^75^. GADD34, downstream of PKR (and PERK) was critical for IFN-β and IL-6 responses to dsRNA and Chikungunya virus^76^. NF-κB and IκB (NF-κB inhibitor) have disparate protein half-lives, thus PERK also contributes to IFNβ mRNA transcription during translation inhibition via the relative decrease in cytosolic ikB^77^. Our study adds to this roster of mechanisms implicating the PERK pathway in IFN control through the suggestion that PERK regulates mitochondrial release of dsDNA during ER stress. In a previous study, PERK deficiency in murine embryonic fibroblasts (MEFs) did not hinder synergism between LPS and ER stress^27^. This may have been a cell type issue, as in the current study a PERK inhibitor decreased both dsDNA release and IFN-β mRNA induction in response to both virus and TPG in HeLa and A549 epithelial cells.

Although there are multiple ways in which PERK could affect mitochondria, we focused on a proximal regulator of mitochondrial permeability. During classical apoptosis, the Bak-Bax pores in mitochondria allow egress of cytochrome c^78^. These pores have also been implicated in dsDNA release^53,55^. Although ER stress does not regulate Bak/Bax directly, it does regulate upstream Bim, increasing Bim mRNA expression by ∼2-fold (confirmed in this study) and protein by ∼5 fold^52^. The CHOP transcription factor, induced by PERK is essential both for Bim upregulation as well as for ER stress-induced apoptosis in macrophages and tunicamycin-induced renal injury in vivo^52^. In these same settings, Bim is also required for ER stress induced apoptosis. In the current study, CHOP deficiency negatively impacted VSV-induced IFN-β. Furthermore, Bim deficiency significantly decreased ER stress-induced dsDNA release in macrophages and was absolutely essential for TPG-stimulated IFN-β mRNA induction. Interestingly, the cells in these studies appeared healthy, even with abundant dsDNA in their cytoplasm and the implication of pro-apoptotic molecules, suggesting mitochondrial dsDNA release may be uncoupled from apoptosis induction, or at least significantly precede apoptosis. Of note, the time frames for the ER stress experiments was short (2-3 hours), whereas continued lower dose TPG exposure has been reported to induce apoptosis in macrophages after 48 hours^79^.

Finally, it was essential to explore the relevance of XBP1 and PERK pathway activation outside of pharmacologic ER stress induction or laboratory oxygen glucose deprivation. Viral infections trigger and modulate the UPR^20,63,80^. Moreover, an increasing literature has identified RNA virus-STING interaction: multiple RNA viruses have evolved mechanisms for targeting both cGAS and STING, including Dengue virus, Zika virus, West Nile virus and Japanese encephalitis virus (recently reviewed in ^59^). The list of RNA viruses that stimulate mtDNA release is growing, suggesting that this may be a general property of RNA viral infection: Dengue virus directly targets mitochondria causing perturbations in structure and function^56,81^. Influenza virus and encephalomyocarditis virus cause mtDNA leakage via M2 and 2B proteins, respectively^43^. Chikungunya virus generates cytosolic DNA^82^. Recently, SARS-CoV2 has also been suggested to trigger mtDNA cytosolic release via viral proteins NSP4 and ORF9b^57,83^. Thus, RNA viral infection appeared a good opportunity to assess the roles and interactions of the UPR, dsDNA release and STING.

Inhibition of either XBP1 or PERK pathway significantly decreased VSV-induced *IFNB1* expression. We had previously described direct binding of XBP1 to an enhancer element 7kb away from the *Ifnb1* gene^50^. Thus, there are other explanations for how XBP1 and PERK might be regulating IFN-β mRNA transcription. However, this study suggests an additional mechanism, in that both XBP1 and PERK contribute to stress-mediated dsDNA release in response to infection with the RNA VSV virus. Induction of the IFN-β gene is highly cooperative, possibly underlying the capacity of multiple mechanisms or agonists to synergize in promoting expression^84^.

The dsDNA release by most of the cells in the culture, even those without GFP evidence of viral infection was surprising. Two possibilities are that these neighboring cells either had low level viral infection (insufficient for visible GFP) or that the infected cells were communicating ER stress (or another message) to uninfected neighbors to begin rallying anti-viral responses. So called “transmissible ER stress” has been described in the cancer literature, in which cancerous cells are able to induce ER stress in invading innate immune cells. Proposed mechanisms include communication via extracellular vesicles, specific protein secretion or metabolites (e.g. lactic acid)^85^. More recently ER stress has been documented to transmit between hepatocytes via CX43 gap junctions^86^. Interestingly, influenza virus-dependent STING signals have also been noted to propagate intercellularly by CX43 gap junctions^43^. Whether these gap junctions or other mechanisms described in the literature play a role in dsDNA release from neighboring cells remains to be determined.

It is not clear whether the UPR inhibition directly impacts MAVS or STING dependent signaling. Although we confirmed a critical role for MAVS in VSV-stimulated IFN production, there was residual IFN induction in *MAVS*^-/-^ cells and STING deficiency decreased IFN induction by over 50%. Furthermore, consistent with previous reports, STING deficiency permitted increased VSV replication^8,58^. In the setting of interferonopathies or other conditions where cytosolic DNA clearance is impaired, STING indirect recognition of mitochondrial DNA might play an even more prominent role, beyond stimulation of MAVs-depenent signaling^44,87^. The effect of UPR inhibition in the absence of MAVS would suggest at least some MAVS ©. However, given multiple cross-talk mechanisms between STING and RIG-1/MAVS signaling, this question is difficult to tease apart further^21,60^.

Together, our results demonstrate that the activation of multiple UPR pathways during certain types of ER stress and viral infections contributes to the generation of cytosolic dsDNA. Ultimately, ER stress induced dsDNA release and subsequent STING activation may be critical for marshaling innate immune responses. On the other hand, in the setting of non-infectious processes generating ER stress, STING may contribute to tissue damage. ER stress-dependent STING activation also potentially alters neoplastic responses. Moving forwards, it will be important to determine the role of the ER stress-STING axis in these disease processes as ER stress is becoming an increasingly modifiable factor^36,88-90^.

## Funding acknowledgements

TH was supported by a UW-Madison Hilldale Scholarship. CRK was an Open Philanthropy Fellow of the Life Sciences Research Foundation. AM is a Burroughs Wellcome Fund Investigator in the Pathogenesis of Infectious Disease and an H.I. Romnes Faculty Fellow funded by the Wisconsin Alumni Research Foundation. This work was supported by National Institutes of Health grants AI164690 to AM.

