## Supplemental figures for "Multiple Unfolded Protein Response pathways cooperate to link cytosolic dsDNA release to Stimulator of Interferon Gene (STING) activation"

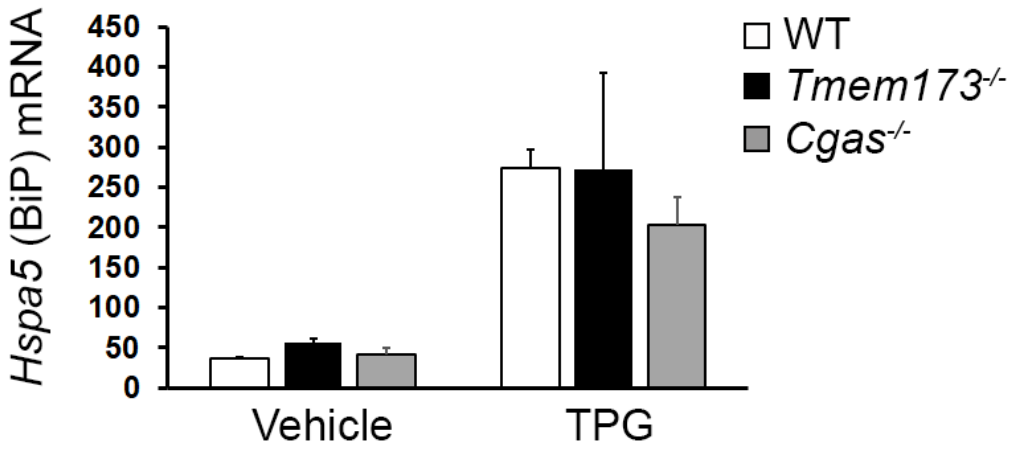


**Figure S1: Thapsigargin induces *Hspa5* (BiP) expression in STING and cGAS knockout macrophages.** Wild type (WT), *Tmem173*-/- or *Cgas*-/- immortalized macrophages were treated with thapsigargin (TPG) for 3h and then harvested for RNA. *Hspa5* expression was detected using qPCR with normalization to 18S rRNA. Bars are from 2 independent experiments with SEM error bars.

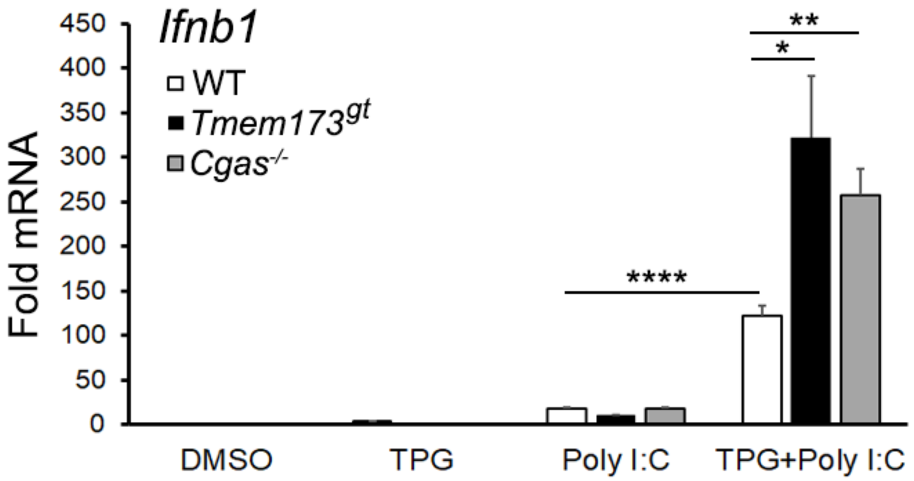


**Figure S2: cGAS and STING are not required for synergistic IFN-β induction between TPG and poly I:C**. WT (white bars), STING null mutant (Golden Ticket, *Tmem173*$^{gt}$ , black bars) or *Cgas*$^{-/-}$ macrophages were stimulated with 1μM TPG for 1h and then 100 μg/mL poly I:C for 6h prior to harvest for RNA. RNA levels were normalized to 18S RNA. Bars represent means of 2-4 independent experiments and errors are SEM.

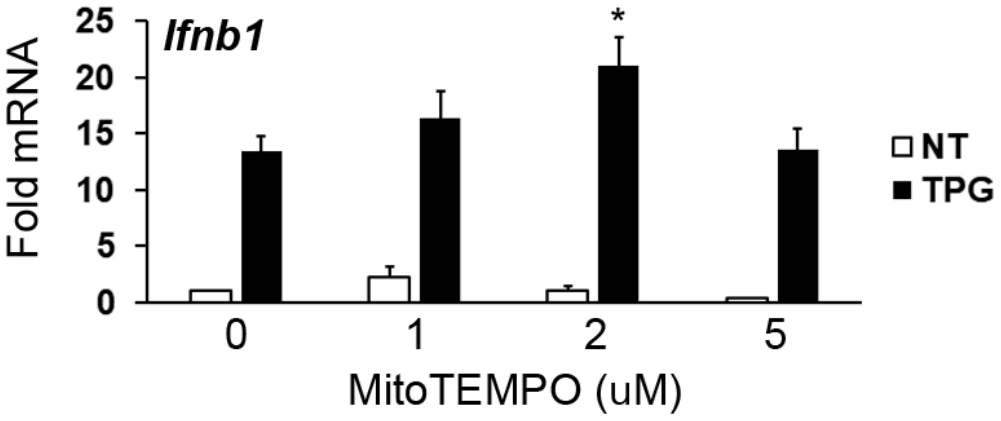


**Figure S3: MitoTEMPO does not decrease Thapsigargin-induced IFN-β expression.** Immortalized macrophages were treated 1h with varying concentrations of MitoTEMPO, the mitochondrial superoxide scavenger, followed by 3h 1 μM Thapsigargin. RNA levels (Fold RNA) were quantitated using qPCR with normalization to 18S rRNA and DMSO vehicle treated (NT) control (set=1). Bars represent means and SEM of 2-6 independent experiments. *$p<0.05$ in comparison with TPG treated control in the absence of MitoTEMPO.

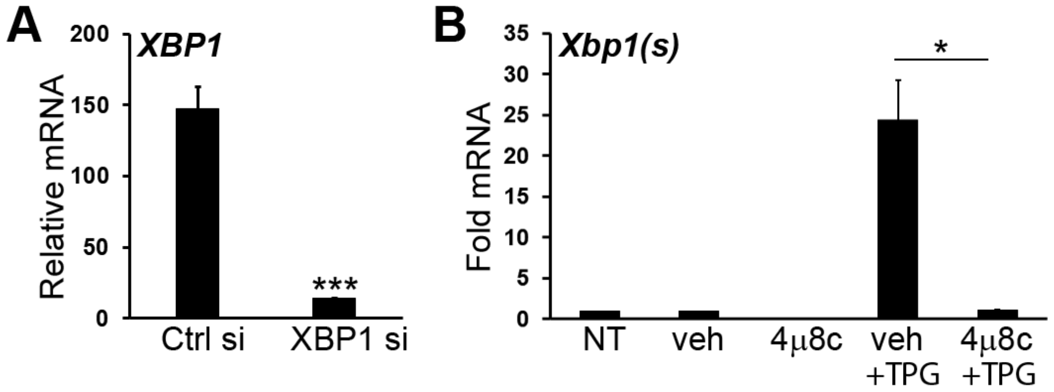


**Figure S4: XBP1 knockdown and 4μ8c inhibition of XBP1 mRNA splicing**. A) HeLa cells were transfected with control (Ctrl) or XBP1 siRNA. After 24h, mRNA was quantitated by qPCR with normalization to 18S rRNA. Bars represent means and SEM of 3 independent experiments. XB1 siRNA knocks down mRNA expression by >90%, ***p=0.0023 vs control siRNA. B) iMac cells were not treated (NT), pre-treated with DMSO (veh) or 10 μM 4μ8c followed by 1 μM TPG for 3h. *p<0.05. Spliced *Xbp1* mRNA was detected by qPCR with normalization to 18S and to NT control (set=1). Results are from 1 experiment in duplicate (SD error bars) and representative of N=3. Similar results were obtained in HeLa and A549 cells.

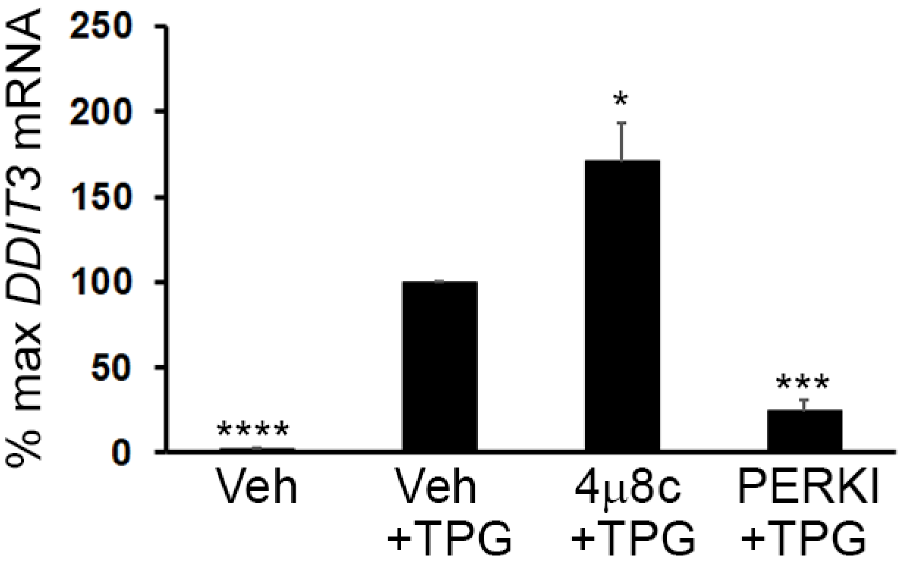


**Figure S5: The PERK inhibitor GSK2606414, but not 4μ8c, blocks TPG-dependent CHOP (*DDIT3*) upregulation**. HeLa cells were pretreated with DMSO vehicle (Veh), the IRE1 inhibitor 4μ8c, or the PERK inhibitor (GSK2606414, PERKI) for 30 minutes, followed by thapsigargin (TPG) for 3 hours. RNA was harvested and cDNA quantitated by qPCR by normalization to 18S rRNA and then to vehicle + TPG (set=100%). Results are from 3 independent experiments performed in duplicate. P-values are vs. vehicle + TPG, * $p<0.05$, ***$p<0.005$, ****$p<0.001$.

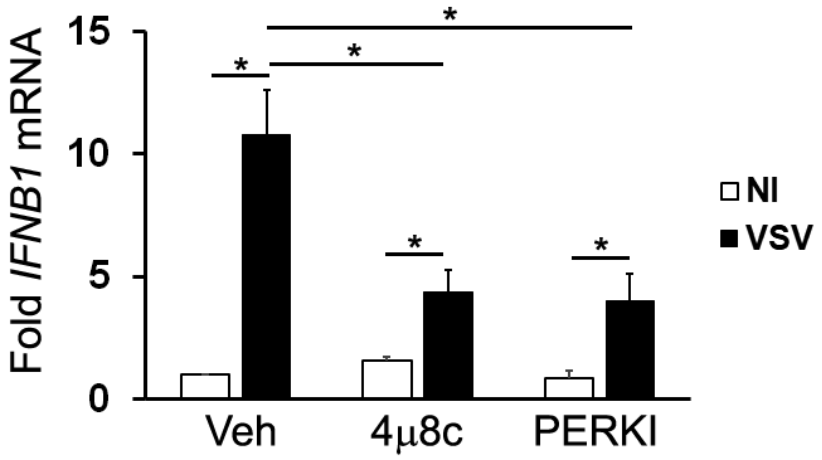


**Figure S6: IRE1 and PERK inhibitors decrease VSV-induced IFN-β expression in HeLa cells**. HeLa cells were pre-treated with DMSO vehicle control or the IRE1 inhibitor 4μ8c or a PERK inhibitor (PERKI) for 30 minutes followed by 6 hours VSV infection. IFN-β mRNA was quantitated by qPCR with normalization to 18SrRNA and to vehicle treated uninfected (NI) control. Results are from 3 independent experiments with SEM, *$p<0.05$ in pairwise comparisons.

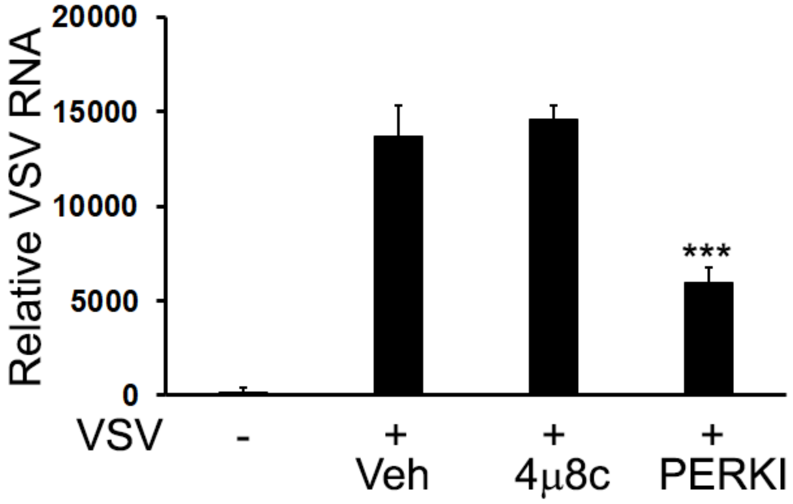


**Figure S7: Effect of UPR inhibitors on VSV replication**. A549 cells were pre-treated for 30 minutes with DMSO vehicle, 4μ8c, or PERK inhibitor (PERKI) and then infected with an MOI=1 of VSV for 24h. VSV replication was detected using qPCR for genomic VSV with normalization to host 18S rRNA. Results are from an experiment in triplicate with SD, and representative of 2 independent experiments. ***$p<0.005$ vs. vehicle+VSV.
